# Homeostatic versus pathological functions of Dual Leucine Zipper Kinase in the adult mouse brain

**DOI:** 10.1101/479378

**Authors:** Sunil Goodwani, Mary E Hamby, Virginie Buggia-Prevot, Paul Acton, Celia Fernandez, Rami Al-Ouran, Yongying Jiang, Michael Soth, Philip Jones, William J. Ray

## Abstract

Dual Leucine Zipper Kinase (DLK, *Map3k12*), is an injury-induced axonal protein that governs the balance between degeneration and regeneration through its downstream effectors c-jun N-terminal kinase (JNK) and phosphorylated c-jun (p-c-Jun). DLK is generally considered to be inactive in healthy neurons until induced by injury. However we report that DLK in the cerebellum appears constitutively active and drives nuclear p-c-Jun in cerebellar granule neurons in the absence of injury. In contrast the adult hippocampus expresses similar levels of apparently constitutively active DLK, but p-c-Jun is lower and does not accumulate in the nucleus. Injury is required there for p-c-jun nuclear expression, because in the rTg4510 model of tauopathy, where there is extensive hippocampal pathology, nuclear p-c-Jun is induced in a DLK-dependent manner. This context-specific regulation of DLK signaling could relate to availability of JNK scaffolding proteins, as the cerebellum preferentially expresses JNK-interacting protein-1 (JIP-1) whereas the hippocampus contains more JIP-3 and Plenty of SH3 (POSH). To understand how DLK signaling differs between the hippocampus and cerebellum, we selectively blocked DLK and measured changes in protein and mRNA expression. In the cerebellum, p-c-Jun levels correlated with synaptophysin, suggesting a link between DLK activity and synaptic maintenance. In rTg4510 mice, hippocampal p-c-Jun instead correlated with markers of neuronal injury and gliosis (Iba1 and GFAP). RNA sequencing revealed that in both brain regions DLK inhibition reduced expression of JNK/c-Jun pathway components and a novel set of co-regulated genes. In the cerebellum, *Jun* mRNA levels were co-regulated with genes mapping to metabolic pathways, while in the rTg4510 hippocampus, *Jun*-correlated mRNAs correspond primarily to neuroinflammation. These data suggest that in the uninjured cerebellum, DLK/p-c-Jun signaling is linked to synaptic regulation, but in the hippocampus, pathologically activated DLK/p-c-Jun signaling regulates genes associated with the injury response.

## Introduction

Neurons, being post-mitotic cells with limited regenerative capabilities particularly in the mammalian central nervous system (CNS), are equipped with sensitive injury surveillance and response mechanisms that ensure that surmountable damage is not lethal, but that irreversible lesion leads to the regulated elimination of cellular structures or even the entire neuron. One of the best characterized injury responses is Wallerian degeneration, the systematic elimination of traumatized axons, during which the distal end of the axon degrades while the proximal end is either preserved for regeneration or transmits apoptotic signals to the cell body (1, 2). DLK, a mixed lineage kinase in the mitogen activated protein kinase (MAPK) pathway is required for this process across species (3). Whereas axotomy or axonal crush leads to profuse neuronal death in a number of experimental settings, in animals lacking DLK, apoptosis is delayed or prevented altogether; in other contexts where regeneration is possible, DLK is required for the re-extension of severed axons (4-9).

In addition to its well-studied role in injury response, DLK is critical for nervous system development as it regulates neuronal migration (10), axon specification and neurite extension (10-12), axon elimination, and programmed cell death (13). Few studies have examined the role of DLK in mature uninjured neurons, particularly in the mammalian CNS. In *Drosophila* and *C. elegans*, the DLK orthologs wallenda (*Wnd*) and DLK-1 regulate presynaptic function and morphology at the neuromuscular junction (14) and limit synaptic strength (15). However, adult somatic knockout of DLK in mice revealed no CNS phenotype other than resistance to various neuronal insults and a modest increase in post-synaptic evoked potentials in the hippocampus (16).

The mechanisms by which DLK subserves distinct roles in degeneration, regeneration, and development are not clear. To perform its function DLK activation must be permissive, as it responds to a wide array of injuries, including excitotoxicity, chemotoxicity, mechanical damage, and neurodegeneration due to pathologically misfolded and aggregated proteins (1, 17). However given that DLK activation can lead to axon retraction and programmed cell death, strict control over its signaling is essential. There are several possible means by which DLK signaling is directed appropriately to the desired context-dependent outcome. One is the subcellular localization of DLK and its JNK effectors. DLK is generally restricted to axons, growth cones, and post-synaptic densities, spatially restricting it from the cell body (8, 18, 19). This compartmentalization likely involves the palmitoylation of DLK, which leads to physical association with transport vesicles (20). A second mechanism is the integrity of axonal transport. Impaired axonal trafficking caused by cytoskeletal disruption or kinesin mutations causes the accumulation of presynaptic cargo in the axon initial segment and cell body and triggers DLK-mediated injury response through an as-yet undescribed mechanism (21, 22). Without disrupted trafficking DLK does not evoke an injury response in these contexts.

The abundance of DLK is also a critical factor in determining outcome. DLK is constrained by rapid proteolysis resulting from ubiquitinylation by Pam/Highwire/RPM-1 (PHR) family members (12, 18, 23-25). Injury-dependent signaling by three redundant Ste20 kinases, MAP4K4, misshapen-like kinase 1 (MINK1/*Map4k6*) and Traf2-and Nck-interacting kinase (TNIK/*Map4k7*) promote DLK stabilization (26). Reducing DLK degradation is sufficient to trigger signaling, since blocking DLK proteolysis or overwhelming degradation by overexpression (27) are sufficient to cause pathway activation, perhaps by generating sufficient protein concentrations to drive homodimerization and intermolecular autophosphorylation (28).

DLK activity is also directed to an appropriate outcome by signalsome assembly, when activated DLK interacts with its direct substrates MKK4/*Map2k4* or MKK7/*Map2k7*, which genetic deletion studies prove are functionally distinct (29), and one of the three JNK isoforms, each of which has unique signaling properties (30). These complexes can be scaffolded onto one of the JIP proteins (JIP-1, JIP-2, JIP-3, POSH/*Sh3rf1* and JLP/*Spag9*) (31-34), which are attached to motor proteins such as kinesins and provide both a platform for signaling crosstalk and a mechanism to transmit signal to the nucleus by retrograde transport. JIP proteins have complex, partially overlapping binding specificities and are substrates for a number of kinases that regulate their affinity for DLK, JNK, and kinesins.

The ultimate output of the signaling cascade is typically induction of AP-1 transcription factors, principally c-Jun, which JNK phosphorylates on S63 or S73, licensing its nuclear translocation and transactivation of apoptotic or pro-regenerative genes (35). This differential response to p-c-Jun reflects in part the availability of nuclear heterodimerization partners (36), highlighting the importance of other signaling pathways in determining the ultimate outcome of DLK activation.

Here, we observed constitutive DLK activity leading to nuclear p-c-Jun accumulation in the adult mouse cerebellum in the absence of injury, suggesting that there must be additional regulatory mechanisms operating to promote proper DLK signaling in mature post-mitotic neurons. We compared DLK-dependent gene expression signatures in the cerebellum, where DLK is constitutively active, to those in the hippocampus, where DLK signaling is restrained in the absence of neuronal insult. We find that DLK signaling elicits different downstream pathways depending on whether its function is homeostatic or injury-induced, perhaps reflecting the differential availability of scaffolding and effector proteins.

## Materials and Methods

### Study design

C57Bl/6 male mice (3-4 months old) used in this study were obtained from the Jackson Laboratory. For all experiments DLKi was formulated in 0.5% methylcellulose and administered by oral gavage (10 mL/kg). Experiments utilizing DLKi were terminated 105 minutes following the last dose (corresponding to the time of maximal plasma and brain drug concentrations) and tissues were isolated as described below. For DLK antibody validation heterozygous matings of B6;129P2-MAP3k12Gt were established to produce wild-type (*Map3k12*+/+), heterozygous (*Map3k12*+/-) and homozygous (*Map3k12*-/-) offspring. Brain tissue was extracted from E16-E20 embryos. rTg4510 mice were obtained from the Jackson Laboratory. All mice were initially group-housed at 22°C with a 12-h light/dark cycle (lights on at 6 am). Food and water were available ad libitum. Prior to experiments, mice were single-housed and randomly assigned to their respective treatment groups that were matched for age, sex, and littermate controls. All experimental procedures were performed according to the National Institute of Health Guidelines for the Care and Use of Laboratory Animals and were approved by the Institutional Animal Care and Use Committee of the University of Texas MD Anderson Cancer Center.

### Tissue harvesting

Once the mouse was deeply anesthetized with 2% isoflurane, the thoracic cavity was opened and the blood was collected using a 1 mL syringe pre-coated with heparin (Sagent; 5,000 Units/ml) and quickly transferred to an EDTA coated Capiject (Terumo) collection tube and mixed for 10 min. Isolated plasma was immediately flash frozen until used for pharmacokinetic analysis. For the mice used in all experiments, except for ActivX KiNativ kinase profiling experiment, a cardiac perfusion was performed following blood collection using 2X PhosSTOP phosphatase inhibitors (Roche) and 1X Complete protease inhibitors (Roche) in 1X PBS pH 7.4 (Ambion). The brain was removed and the hemispheres were separated at the midline. The right hemisphere was further dissected to isolate cerebellum, brainstem, hippocampus and remaining forebrain and snap frozen. For all experiments except for RNASeq experiment, left hemisphere was postfixed in 4% paraformaldehyde in PBS (Affymetrix) and kept at 4ᵒC overnight. The left hemisphere was then sequentially cryoprotected in 20% sucrose followed by 30% sucrose and maitained at 4ᵒC. The cryoprotected hemisphere was then sectioned sagitally at 30 µm using a cryostat (Leica Biosciences). Sections were preserved in cryoprotectant (30% glycerol, 30 ethylene glycol and 1xPBS) and stored at −20ᵒC. For RNAseq experiment left hemisphere was dissected to isolate cerebellum, brainstem, hippocampus and remaining forebrain in Eppendorf™ Snap-Cap Microcentrifuge Biopur™ Safe-Lock™ Tubes and snap frozen for RNA isolation.

### Antibodies and DLKi

The DLKi used in this study was selected by screening published DLK inhibitors from patent and scientific literature and corresponds to example 68 in patent WO 2015/091889. The primary antibodies used for immunoblot analysis were as follows: DLK, 1:250 (NeuroMab; clone N377/20, 75-355); DLK, 1:250 (Invitrogen, PA532173); p-c-Jun, 1:100 (Cell Signaling Technology, 9261); c-Jun, 1:250 (Cell Signaling Technology, 9165); GAPDH, 1:5000 (Millipore, MAB374); phospho-PHF-Tau Ser202/Thr205, 1:1000 (Life Technolgies, MN1020); PSD95, 1:500 (Cell Signaling Technologies, 2507); Synaptophysin, 1:500 (Cell Signaling Technologies, 4329); GFAP, 1:500 (Cell Signaling Technologies, 3670); Iba1, 1:250 (Wako, 01620001); p-MKK4 Ser57/Thr261, 1:100 (Cell Signaling Technologies, 9156); MKK4, 1:200 (Cell Signaling Technologies, 9152); MKK7, 1:200 (Cell Signaling Technologies, 4172); JIP1, 1:250 (Abcam, Ab24449), JIP-2, 1:250 (Abcam, Ab154090); Jip-3, 1:200 (Life Technologies, PA536369); POSH, 1:500 (Proteintech, 14649-1-AP); JNK1, 1:500 (Cell Signaling Technologies, 3708); JNK2, 1:500 (Cell Signaling Technologies, 9258); JNK3, 1:500 (Cell Signaling Technologies, 2305); p-JNK Thr183/Tyr185, 1:500 (Cell Signaling Technologies, 9251), JNK, 1:500 (Cell Signaling Technologies, 9252). Secondary antibodies used for immunoblot detection were Goat anti-rabbit, 1:5000 (Licor, Catalog number 926-68021, Fluorophore: IRDye 680LT); Goat anti-mouse, 1:5000 (Licor, Catalog number 926-68020, Fluorophore: IRDye 680LT). Primary antibodies used for immunofluorescence and immunohistochemistry staining were as follows: DLK (Invitrogen), and phospho c-Jun (Cell Signaling). Donkey anti-rabbit and donkey anti-mouse Alexafluor-conjugated secondary antibodies were used for fluorescence microscopy (Life Technologies).

### Tissue processing for Western blot analysis

Cerebellar, hippocampal and frontal cortical tissues isolated the mice were lysed in approximately 10 volumes of RIPA lysis buffer (150 mM NaCl, 1.0% IGEPAL^®^ CA-630, 0.5% sodium deoxycholate, 0.1% SDS, 50 mM Tris, pH 8.0) containing 2X Halt Protease inhibitors (Thermo Scientific) and 2X Halt phosphatase inhibitors (Thermo Scientific) by mechanical homogenization on ice until a homogenous suspension was obtained. Lysates were centrifuged at 14,000Xg at 4ᵒC for 30 minutes. The supernatants were snap frozen on dry ice and stored at −80ᵒC. Total protein concentration was determined using DC Protein Assay (Bio-Rad). Lysates were diluted with 4X protein sample loading buffer (Licor) containing 10% 2-mercaptoethanol and heated to 95ᵒC for 10 min. The samples were then subjected to gel electrophoresis on NuPAGE 4-12% Bis-Tris Gels (Life technologies) using 1X MES running buffer (Life Technologies) after which they were transferred to nitrocellulose membranes (Life Technologies) and incubated with Odessey Blocking Buffer (Licor) for 1 h at room temperature. The membranes were then incubated with primary antibodies overnight at 4ᵒC with continual shaking, washed three times with 1X TBS-0.1% Tween (TBST) at room temperature with continuous shaking, and then incubated in secondary antibodies for 1 h at room temperature with continuous shaking before washing with TBST three times. The membranes were then imaged using the Licor Odyssey Imager and quantified using Image Studio 4.0 Software (Licor). The data were analyzed using Microsoft Excel and GraphPad Prism.

### Immunohistochemistry and immunofluorescence analysis

Free floating sagittal brain sections (30 µm) were washed in TBS and subjected to antigen retrieval for 20 min at 90°C in a sodium citrate buffer (Sigma). Endogenous fluorescence was quenched by incubation in 10 mM glycine in TBS with 0.25% Triton X-100 (TBS-T) followed by blocking in 5% horse serum in TBS-T. Sections were incubated overnight at 4°C in primary antibodies diluted in 1% BSA in TBS-T, followed by secondary antibody incubation for 2 h at room temperature. Sections were stained for nuclei using Hoechst 33342 (Life Technologies), mounted on Superfrost Plus slides (Fisherbrand) in VectaShield mounting media (Vector Labs) and allowed to cure overnight before imaging. DAB immunohistochemistry was performed using Vector ABC (Vector Labs) according to the manufacturer’s instructions. After the final reaction was terminated, sections were mounted on Superfrost Plus slides in Cytoseal XYL mounting media (Sigma) and allowed to cure overnight before imaging. Images were acquired on a Nikon Eclipse Ti confocal microscope using 20X and 60X objectives and NIS Elements Imaging Software (Nikon). For comparison between conditions, the same acquisition settings were used for each channel across samples. All images were processed using ImageJ software (NIH), using the signal from control IgG staining to set the background. To assess the level of p-c-Jun immunofluorescence in Hoechst-positive nuclei in the brain, a mask was created in the Hoechst image, applied to the corresponding p-c-Jun image, and fluorescence outside the region defined by the mask was cleared. The remaining fluorescence was quantified by measuring the raw integrated density in the field, which is the sum of the pixel values in the image. The raw integrated density signal was then expressed as a percentage of the signal in the cerebellum of vehicle-treated control.

### RNASeq Analysis

Snap frozen tissue (cerebellum and hippocampus) was homogenized per manufacturer’s recommendation using Qiagen RNeasy Mini kit lysis buffer and and Qiagen Tissuerupter and RNA purified using the Qiagen RNeasy Mini kit. mRNA-seq was performed using Illumina HiSeq 4000 with 76 paired-end reads. Sequence read quality was assessed using FastQC v0.11.5 (59) before and after adapter trimming. Adapter trimming was conducted using Cutadapt v1.15 (60). The trimmed paired end read sequences were then aligned to the mouse reference genome GRCm38 (mm10) using the STAR aligner v2.5.1b (37), (38). Genes were quantified using the STAR aligner option quantMode which utilizes the HTSeq algorithm (39). Low counts were discarded and data was normalized (standard method), followed by differential gene expression analysis (ANOVA) using Partek^®^ Flow^®^ version 7.0 (Copyright 2018; Partek Inc.). Heatmap was generated using Heatmapper (40). Ingenuity Pathway Analysis was performed as previously described (41). DiRE was used for promoter analysis (42).

STRING Analysis (version 10.5) was used to examine the most highly interconnected genes correlated with Jun (43).

### Pharmacokinetic assessment of DLKi in plasma and brain tissue

Concentration of DLKi in both plasma and brain tissue samples was quantitated by LC-MS/MS after sample processing. For plasma sample analysis, 25 μL of each sample was precipitated with 200 μL of acetonitrile containing 5 ng/mL of a reference compound that served as the internal standard (IS). This suspension was vortexed for 30 min and centrifuged at 4k rpm for 15 min, after which 100 μL of the extract was aliquoted and diluted with 200 μL of water mix prior to LC-MS/MS analysis. For brainstem tissue sample analysis, the tissue samples were homogenized in 80% methanol/water (1 mL per 100 mg tissue) at 4 °C using OMNI Bead Ruptor 24 coupled with Omini BR-Cryo cooling unit (Omni International, Kennesaw, GA). After centrifugation at 15k rpm for 15 min, the supernatant (60 μL) was mixed with 50% acetonitrile/water (140 µL) containing 5 ng/mL IS prior to LC-MS/MS analysis. LC-MS/MS analysis was conducted on a Waters Acquity UPLC system coupled with a Waters TQ-S triple quadrupole mass spectrometer system operated at positive mode (Waters, Manchester, United Kingdom). The DLKi and IS were separated using a Supelco Ascentis fused-core C18 column (2.7 μm, 2.1 × 20 mm) (Sigma-Aldrich, St. Louis, MO) and detected by multiple reaction monitoring transitions. The mobile phase A was 0.1% acetic acid-water and B was 0.1% acetic acid-acetonitrile. The gradient was 15% B (0-0.3 min), 15-95% B (0.3-1.3 min), 95-15% B (1.3 to 1.31 min), 15% B (1.31 to 1.7 min) and the flow rate was 0.5 mL/min. The column temperature was 40 °C. The injection volume was 2 µL. Under these conditions, the retention time was 0.8 min for DLKi and 1.08 min for IS. The method was validated with analytical range of 1 – 1000 ng/ml DLKi in both untreated mouse plasma and brain tissue homogenate.

### Pharmacokinetic/Pharmacodynamic (PK/PD) Analysis

PK/PD analysis was performed using GraphPad Prism software. The percent change in cerebellar p-c-Jun/c-Jun levels relative to vehicle control obtained from the western blot analysis was utilized as the pharmacodynamic response and plotted against the PK parameter. The four parameter logistic curve was utilized to obtain the IC_50_ values.

## Results

### DLK drives c-Jun phosphorylation in the adult mouse cerebellum

We assessed DLK expression in three regions of the adult C57Bl/6 mouse brain (Fig. 1A). In each region, DLK immunoreactivity broadly labeled neurons and their processes, with particular intensity in the molecular layer of the cerebellum and in the mossy fibers extending to the CA3 region of the hippocampus (Fig. 1B). Morphological examination revealed that DLK is predominantly axonal, consistent with previous reports (44). Next we examined p-c-Jun; intense immunoreactivity was noted in the nuclei in the granule cell layer of the cerebellum within cells that have the morphological appearance of cerebellar granule neurons (Fig. 1D,E). As these animals were handled for 4 days prior to sacrifice to prevent stress-induced p-c-Jun expression, p-c-Jun was sparse in other brain regions; approximately 10-fold lower throughout the cortex and hippocampus as measured by total nuclear fluorescence intensity, with only occasional p-c-Jun-positive nuclei (Fig. 1F).

**Fig. 1.**
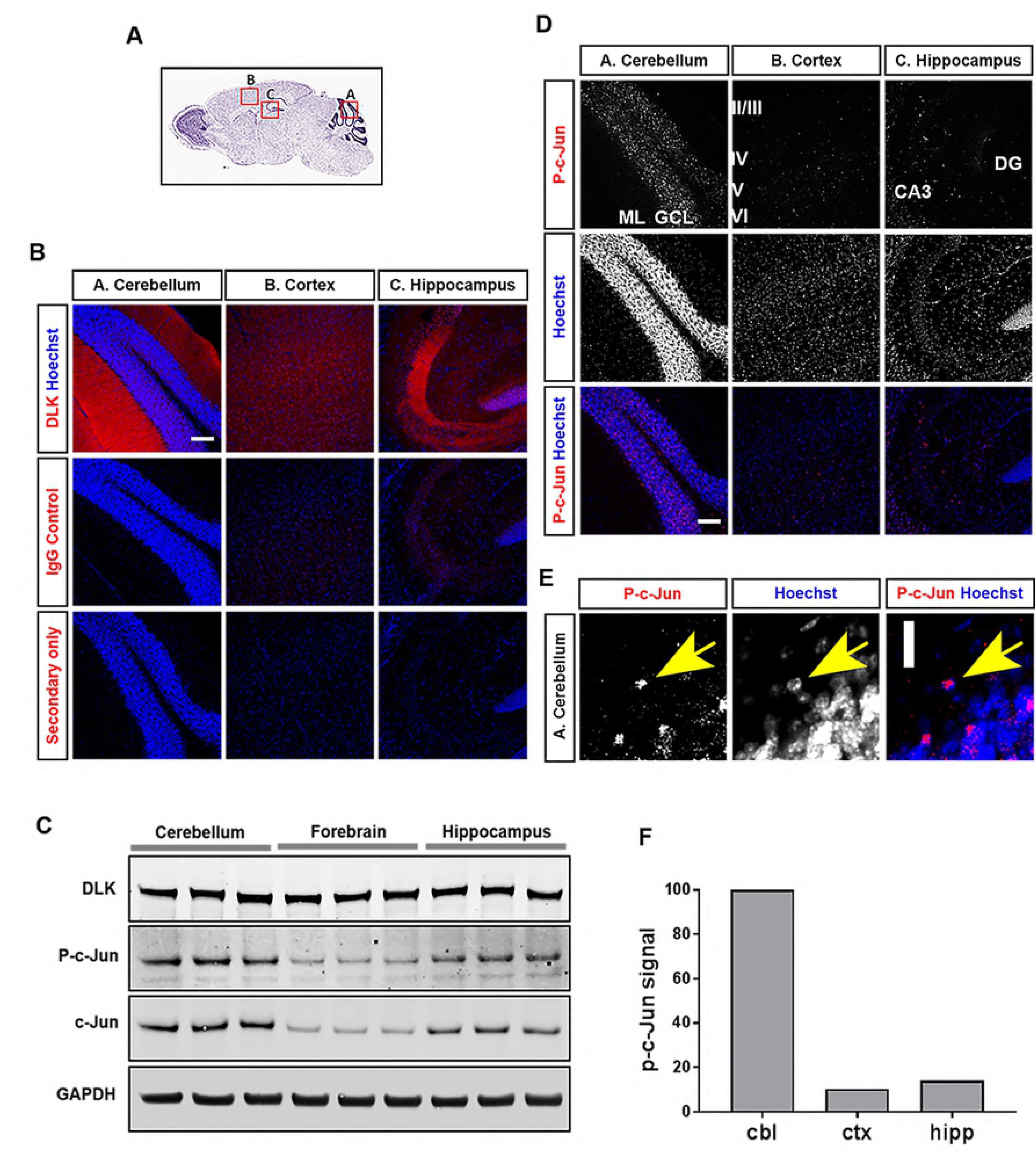
DLK constitutively signals in adult uninjured mouse cerebellum. **(A)** Nissl-stained sagittal section from the Allen Mouse Brain Atlas for reference; red boxes indicate the areas shown in the lower panel. **(B)** Representative double staining for DLK (red) and Hoechst (blue) shows its distribution in the molecular layer of the cerebellum (A), diffusely throughout the cortex (B) and in the mossy fiber axon terminals in the hippocampus (C) of adult uninjured C57Bl/6 mouse brain. Negligible staining is observed with rabbit IgG or secondary alone. Scale bar: 100um. **(C)** Immunoblot analysis of DLK, p-c-Jun and c-Jun in lysates obtained from cerebellum, forebrain and hippocampus of 3-4 months old C57Bl/6 mice. **(D)** Representative double staining for p-c-Jun (red) and Hoechst (blue) shows its distribution overlapping with Hoechst-positive nuclei in the in the granule cell layer of the cerebellum (A), in layer V of the cortex (B) and to a lesser degree in the CA3 region of the hippocampus (C). GCL, granule cell layer; ML, molecular layer; CA3, Cornu Ammonis subfield 3 of the hippocampus; DG, dentate gyrus. Scale bar: 100 μm. **(E)** p-c-Jun (red) fluorescence overlaps with Hoechst (blue)-positive nuclei, indicating a nuclear localization. Scale bar: 20 μm. **(F)** Quantification of p-c-Jun fluorescence signal in Hoechst-positive regions in cerebellum (cbl), cortex (ctx) and hippocampus (hipp) expressed as a percentage of signal in cerebellum.

DLK, p-c-Jun, and c-Jun levels in in forebrain, hippocampus, and cerebellum was also assessed by Western blot. We first confirmed using DLK -/- animals the specificity of two different DLK antibodies, as both detected a band at 120 kD that was absent in the DLK -/- mouse brain lysate (Fig. S1A). Moreover the 120 kD species was verified as mouse DLK by immunoprecipitation followed by mass spectrometry (Fig. S1B). Using these validated reagents, full-length DLK was expressed at similar levels in all three brain regions (Fig. 1C). The biochemical assessment of p-c-Jun aligned with the immunofluorescence data: levels were higher in cerebellum and lower in the hippocampus and forebrain (Fig. 1F). As would be predicted, since p-c-Jun drives its own expression (45), c-Jun levels were also lower in those regions.

Given this result we asked whether the DLK drives the phosphorylation and nuclear accumulation of c-Jun in the cerebellum. To block DLK activity fully throughout the brain, we used a highly selective DLK inhibitor (DLKi) with good oral bioavailability and CNS penetration. Acutely blocking DLK (105 min) rather than using conditional knockout has two advantages: virtually complete DLK inhibition is possible, and the short duration of inhibition helps limit the findings to direct effects of kinase inhibition rather than any potential compensation. We administered DLKi orally to C57Bl/6 mice and assessed cerebellar p-c-Jun levels by Western blot and immunofluorescence. At 105 min post single dose of DLKi, nuclear p-c-Jun immunofluorescence in the granule cell layer was nearly completely abolished in a dose-dependent manner (Fig. 2A-B). Western blots of cerebellar lysates indicated that DLK protein level was unchanged, but p-c-Jun levels declined approximately 3-fold in 10 mg/kg and 30 mg/kg DLKi-treated mice compared to vehicle treated mice (Fig. 2C-D)(P value<0.0001, one-way repeated measures ANOVA, followed by Dunnett’s t-test). Phosphorylated JNK (pJNK) as assessed by Western blot analysis here and in subsequent experiments were not significantly altered by DLKi, however this finding was expected since only a small percentage of the total pJNK pool is DLK-dependent (17). To confirm the selective action of DLKi in vivo, we used KiNativ kinase profiling, a mass-spectrometry-based target engagement method that measures all the kinase(s) a drug has bound in tissue (46). KiNativ profiling of brain lysates harvested 105 min after dosing with DLKi demonstrated that of the 232 kinases detectable in brain homogenate, only DLK was fully occupied, with lower but measurable binding to the DLK homolog leucine zipper kinase (LZK) and Jak1 site 1, demonstrating high selectivity (Fig. 2E). At brain concentrations associated with p-c-Jun reduction (>10 μM), DLK was >95% bound to DLKi in the brain, whereas LZK and JAK1 was bound ~50% or less (Fig. 2F). Increasing the dose to achieve greater inhibition of LZK or other kinases did not further reduce p-c-Jun (Fig. 2D). Thus, we conclude that DLK is required for the majority of the p-c-Jun in the cerebellar granule neurons.

**Fig. 2.**
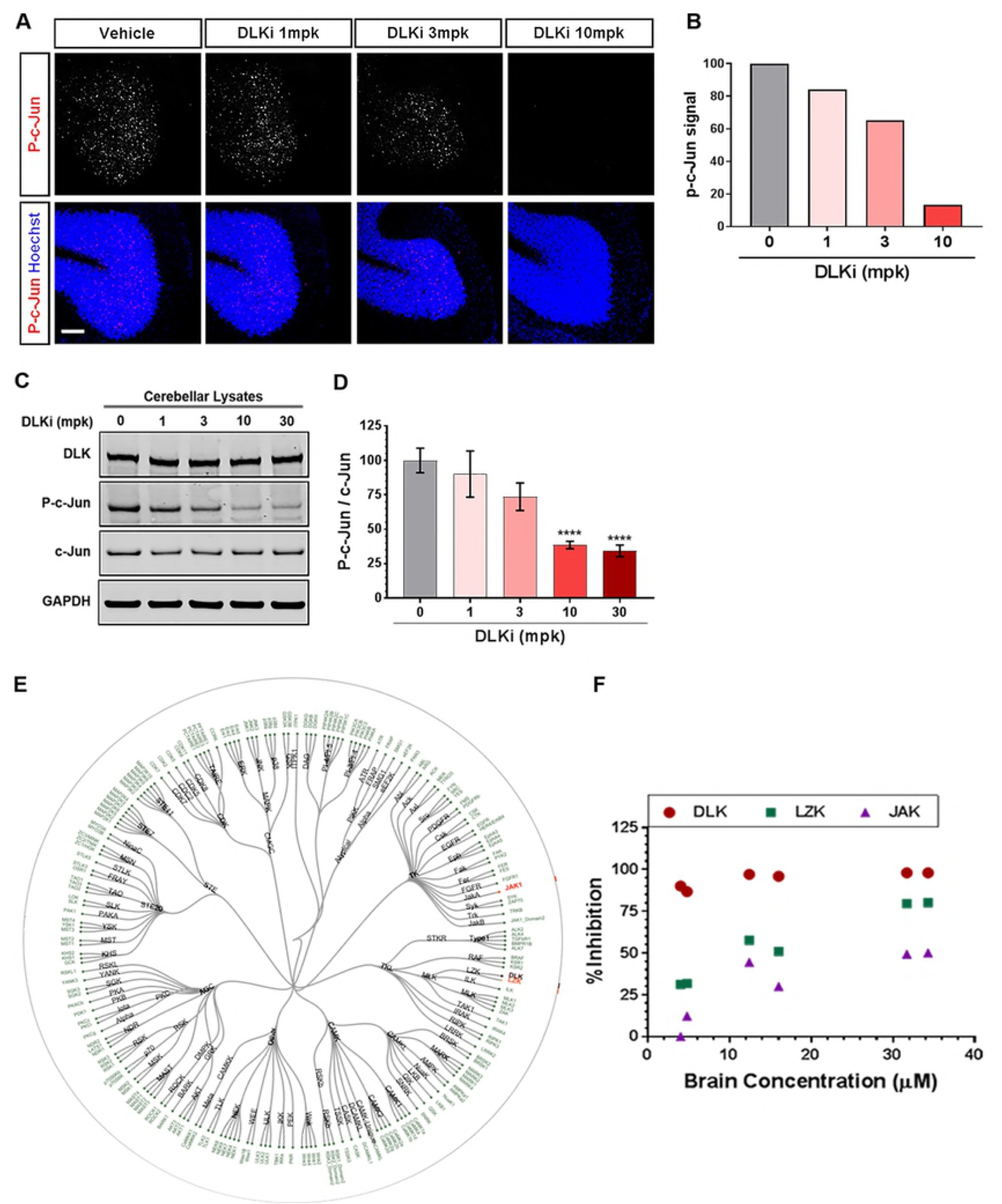
Acute pharmacological inhibition of DLK significantly reduces c-Jun phosphorylation in adult uninjured mouse cerebellum. **(A)** Representative staining for p-c-Jun in the cerebellum of 3-4 months old C57Bl/6 mice dosed with vehicle or DLKi at 1, 3, or 10 mg/kg. **(B)** Quantification of p-c-Jun fluorescence in cerebellar nuclei expressed as a percentage of the signal from vehicle-treated control. Scale bar: 100 μm. **(C)** Representative immunoblot analysis of DLK, p-c-Jun, c-Jun and GAPDH in cerebellar lysates of 3-4 months old C57Bl/6 mice dosed with vehicle (n=7) or DLKi at 1 (n=4), 3 (n=4), 10 (n=8) or 30 (n=8) mg/kg. **(D)** Quantification of p-c-Jun, normalized to c-Jun, in cerebellar lysates of 3-4 months old C57Bl/6 mice dosed with vehicle (n=7) or DLKi at 1 (n=4), 3 (n=4), 10 (n=8) or 30 (n=8) mg/kg, plotted as percent mean fold change relative to vehicle treated group. ****P < 0.0001, One-way repeated measures ANOVA, followed by Dunnett’s test. **(E)** Graphical representation of the kinases examined for kinome selectivity in C57Bl/6 mice dosed with DLKi by ActivX KiNativ analysis. **(F)** Percent inhibition of DLK (red circles), LZK (green squares) and JAK (purple triangles) in 3-4 months old C57Bl/6 mice brain tissue dosed with DLKi at 2mg/kg (n=2), 20mg/kg (n=2) and 60mg/kg (n=2) assessed by ActivX KiNativ assay plotted against the concentration of DLKi in brain in µM.

### DLK is activated in the forebrain of rTg4510 mice by pathological tau

DLK is generally regarded as injury induced, leading us to question how DLK-mediated signaling differs between the cerebellum, where its activation does not cause degeneration, and the hippocampus and forebrain, where its activation contributes to synapse elimination and neuronal loss (17). To study this question we employed rTg4510 mice, which express high levels of P301L mutant human tau specifically in the forebrain and hippocampus, causing widespread neuronal loss in those regions in an age-dependent manner by 6 months of age (Fig. 3A), but sparing the cerebellum (47). A similar mouse model (Tau P301L) was used to demonstrate that DLK is required for nuclear p-c-Jun accumulation and contributes to neuronal loss in the subiculum (17).

**Fig. 3.**
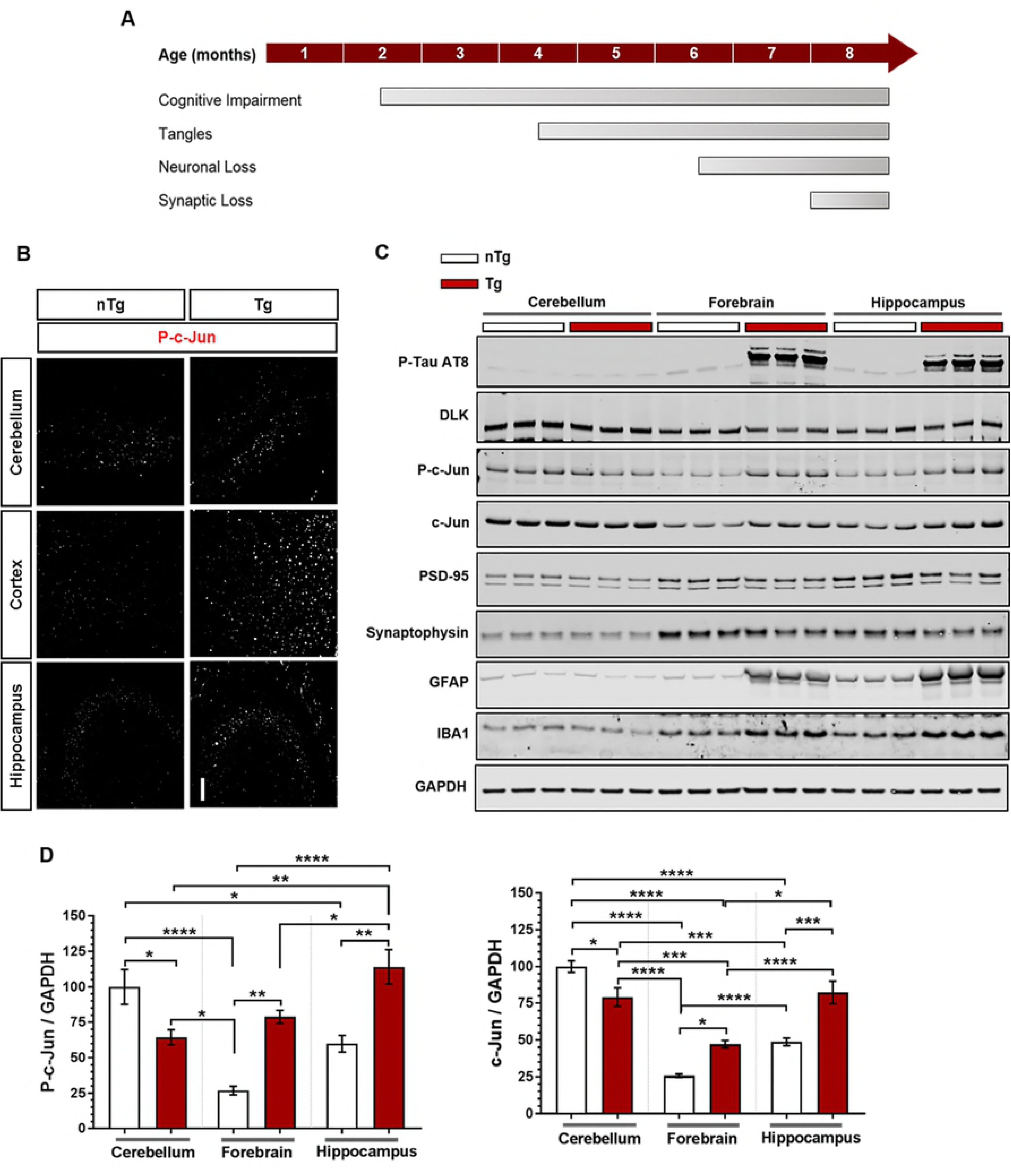
DLK/p-c-Jun pathway is activated in rTg4510 mice forebrain. **(A)** Schema depicting the progression of disease in rTg4510 mice. **(B)** Representative images of p-c-Jun staining in the cerebellum, cortex and hippocampus of 7-8 months old nTg and Tg mice. Scale bar: 100 μm. **(C)** Representative immunoblot analysis of p-Tau AT8, DLK, p-c-Jun, c-Jun, PSD-95, synaptophysin, GFAP and IBA1 from the lysates obtained from cerebellum, cortex and hippocampus of 7-8 months old nTg (white boxes) and Tg (red boxes) mice. **(D)** Quantification of p-c-Jun (n=6), normalized to GAPDH, and c-Jun (n=6), normalized to GAPDH, from the lysates obtained from cerebellum, cortex and hippocampus of 7-8 months old nTg (white bars) and Tg (red bars) mice, plotted as mean percent change relative to nTg cerebellum group. *P < 0.05, **P < 0.01, ***P < 0.001, ****P < 0.0001, One-way repeated measures ANOVA, followed by Tukey’s multiple comparisons test.

Immunofluorescence staining on sagittal brain sections from 7-8 month old rTg4510 (Tg) mice and non-transgenic littermates (nTg) allowed us to compare forebrain, hippocampus, and cerebellum and examine p-c-Jun levels. We observed an elevation of nuclear p-c-Jun in the hippocampus and forebrain of Tg mice relative to nTg mice (Fig. 3B). Western blots with the AT8 antibody recognizing hyperphosphorylated tau confirmed that Tg brain lysates from the forebrain and hippocampus contained high levels of pathological tau (Fig. 3C). As expected cerebellar lysates (from both nTg and Tg) had negligible amounts of hyperphosphorylated tau. In line with our immunofluorescence data, Western blot analysis further confirmed that Tg mice, as compared to nTg, had elevated levels of p-c-Jun in the brain regions (forebrain and hippocampus) with high levels of hyperphosphorylated tau (Fig. 3C-D). Next we examined markers of neuronal injury and synaptic proteins. Not surprisingly the accumulation of hyperphosphorylated tau and elevation of p-c-Jun in forebrain and hippocampus of Tg mice was also associated with increase in the markers of gliosis, GFAP and IBA1, denoting astrocytic and microglial activation, respectively.

Pearson correlation analysis of the relative protein expression of GFAP and IBA1 revealed a significant positive correlation with p-c-Jun in the forebrain and hippocampus (R^2^=0.788, P value<0.0001 for GFAP and R^2^=0.871, P value<0.0001 for IBA1), but no such correlation was found in the cerebellum (R^2^=0.006, P value=0.809 for GFAP and R^2^=0.197, P value=0.149 for IBA1) (Fig 4). In contrast in the cerebellum there was significant correlation between p-c-Jun and synaptophysin, a presynaptic marker, (R^2^=0.896, P value<0.0001) and also to a lesser degree between p-c-Jun and PSD-95, a post-synaptic protein (R^2^=0.333, P value<0.05) (Fig. 4). These data further suggested that DLK function differs depending on whether it is induced by pathological stimuli, in which case p-c-Jun correlates with gliosis, or is constitutively active, in which case p-c-Jun corresponds to synaptic markers.

**Fig. 4.**
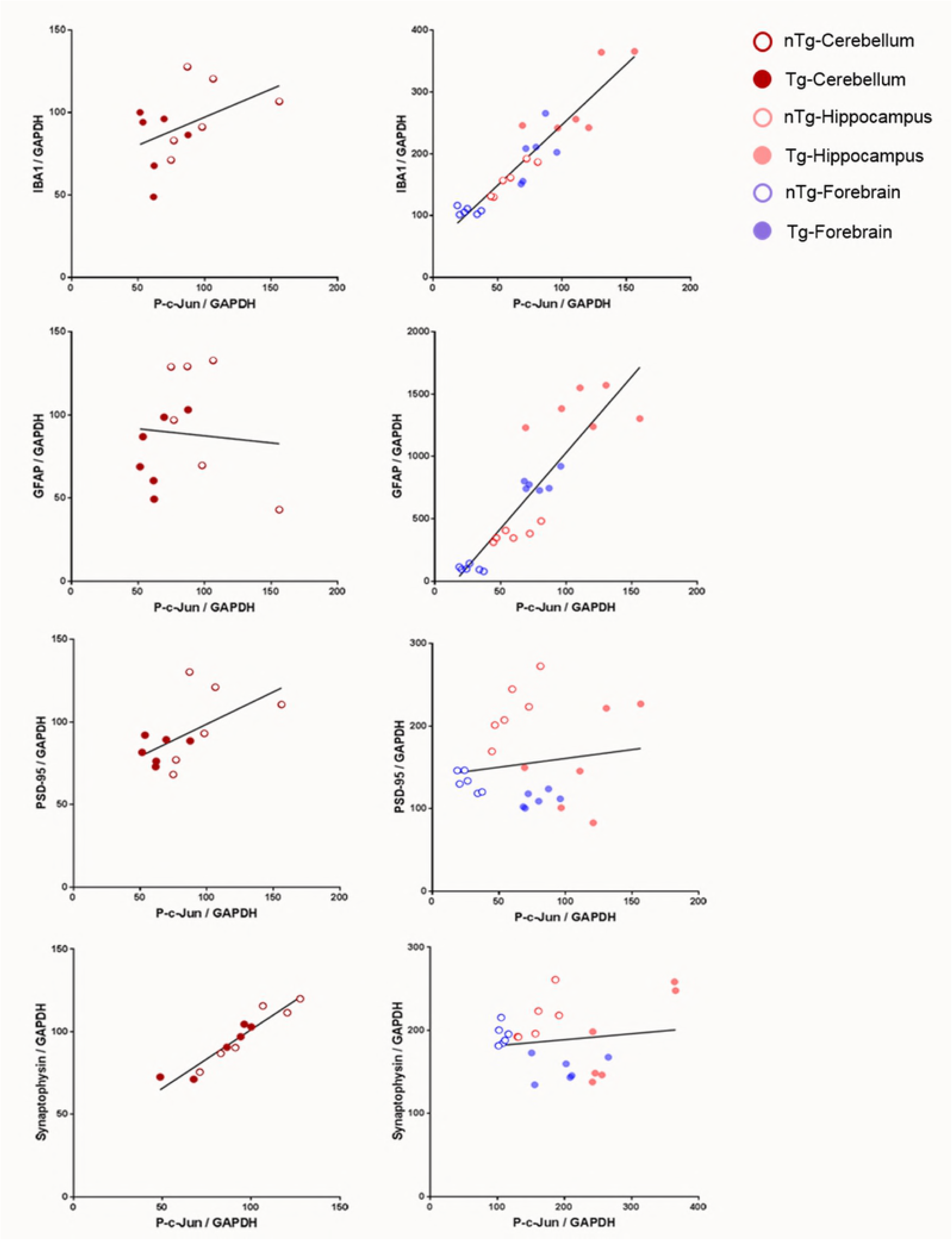
DLK dependent p-c-Jun correlates with different markers in forebrain and cerebellum of rTg4510 mice. Correlation plots between the amount of IBA1, GFAP, PSD-95 and synaptophysin with p-c-Jun in cerebellum, forebrain and hippocampus of 7-8 months old nTg and Tg mice. Open circles, nTg; closed circles, Tg; red, cerebellum, pink, forebrain; purple, hippocampus. Cerebellum Iba1 and p-c-Jun R^2^=0.197, P value=0.149; forebrain and hippocampus Iba1 and p-c-Jun R^2^=0.871, P value<0.0001; cerebellum GFAP and p-c-Jun R^2^=0.006, P value=0.809; forebrain and hippocampus GFAP and p-c-Jun R^2^=0.788, P value<0.0001; cerebellum PSD-95 and p-c-Jun R^2^=0.333, P value=0.049; forebrain and hippocampus PSD95 and p-c-Jun R^2^=0.021, P value=0.498; cerebellum synaptophysin and p-c-Jun R^2^=0.896, P value<0.0001; Forebrain and hippocampus synaptophysin and p-c-Jun R^2^=0.021, P value<0.491; Pearson Correlation.

### DLK drives nuclear p-c-Jun accumulation in the forebrain and hippocampus of rTg4510 mice

We initially tested whether DLK inhibition would reduce the p-c-Jun observed in response to pathological tau by dosing a small cohort of 7-8 month-old rTg4510 mice for 5 days with DLKi (Fig. 5A), followed by immunohistochemistry and Western blot. Five days was chosen to ensure that sufficient time was allowed to capture DLK-dependent p-c-Jun in this disease model. In the nTg mice, as seen in C57Bl/6 mice (Fig. 1), there was abundant nuclear p-c-Jun in the cerebellar granule cells but sparse nuclear p-c-Jun in the cortex and CA3 region of the hippocampus (Fig. 5B). In the Tg mice, as compared to nTg mice, there was marked p-c-Jun immunoreactivity in the nuclei of cortical and CA3 hippocampal neurons, which was nearly eliminated by the DLKi. These observations were further confirmed by Western blot (Fig. 5C). DLK levels remained identical in all brain regions regardless of genotype and treatment. In the cerebellum, p-c-Jun was equally present in both nTg and Tg animals and similarly reduced by DLKi. In contrast, the Tg mice showed increased p-c-Jun in the hippocampus and cortex; this induction was reduced to nTg levels after dosing with the DLKi.

**Fig. 5.**
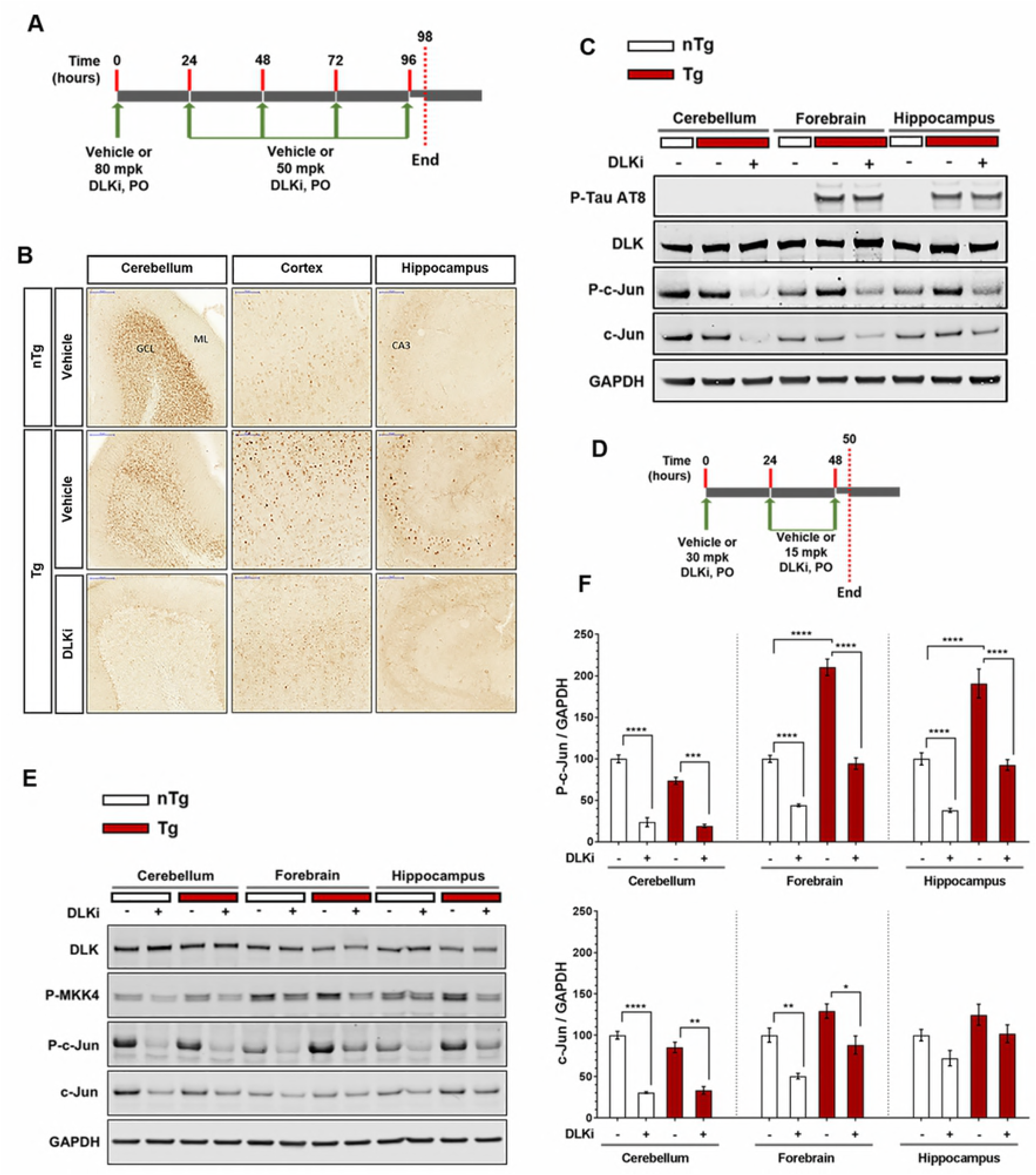
DLKi treatment reduces c-Jun phosphorylation in cerebellum and forebrain of rTg4510 mice. **(A)** Schematic representation of study timeline. nTg (n=3) and Tg (n=3) mice were treated with vehicle or DLKi (80 mg/kg, QD, day 1 and 50 mg/kg, QD, days 2-5). This dosing regimen was chosen based on pharmacokinetic modeling to ensure consistent exposure on all study days; the study was terminated approximately 2 h after the last dose at the maximal plasma concentration of DLKi. **(B)** Representative images of p-c-Jun staining in the cerebellum, cortex and hippocampus of nTg and Tg mice treated with vehicle or DLKi. **(C)** Representative immunoblot analysis of phosho-Tau AT8, DLK, p-c-Jun and c-Jun from the lysates obtained from cerebellum, forebrain and hippocampus of nTg (white boxes) and Tg (red boxes) mice treated with vehicle or DLKi. **(D)** Schematic representation of study timeline. nTg (n=6) and Tg (n=6) mice were treated with vehicle or DLKi (30 mg/kg, qd, day 1 and 15 mg/kg, qd, days 2-3). The study was terminated approximately 2 hours after the last dose. **(E)** Representative immunoblot analysis of DLK, p-MKK4, p-c-Jun and c-Jun from the lysates obtained from cerebellum, forebrain and hippocampus of nTg (white boxes) and Tg (red boxes) mice treated with vehicle or DLKi. **(F)** Quantification of p-c-Jun (n=6), normalized to GAPDH, and c-Jun (n=6), normalized to GAPDH, from the lysates obtained from cerebellum, forebrain and hippocampus of nTg (white bars) and Tg (red bars) mice treated with vehicle or DLKi, plotted as mean percent change relative to nTg vehicle group in respective brain region. *P < 0.05, **P < 0.01, ***P < 0.001, ****P < 0.0001, One-way repeated measures ANOVA, followed by Tukey’s multiple comparisons test.

Having observed DLK dependent p-c-Jun in the Tg mouse forebrain and hippocampus, we repeated this experiment with larger numbers of animals to quantify the data and to examine pMKK4, a direct product of DLK activity. The other known direct produce of DLK, pMKK7, was not measurable reliably in mouse brain with any available antibodies and thus was not included. We chose a shorter duration and lower dose to minimize secondary effects of DLK inhibition or off-target activity (Fig. 5D). Again there was no change in DLK levels in any condition, and DLKi in the cerebellum decreased p-c-Jun 76.4% in nTg and 73.9% in Tg mice (both comparison P value<0.001, one-way repeated measure ANOVA, followed by Tukey’s multiple comparisons test), with no effect of transgenic status (Fig 5F). In the forebrain and hippocampus, In the Tg mice, p-c-Jun was significantly elevated compared to nTg 2.1-fold in forebrain and 1.9-fold in hippocampus (both comparisons P value<0.0001, one-way repeated measure ANOVA, followed by Tukey’s multiple comparisons test); this elevation was abolished by DLKi. p-c-Jun was reduced by DLKi by 55.9% and 55.1% in the nTg and Tg forebrain respectively, and 62.1% and 51.4% in the nTg and Tg hippocampus, respectively (all of these comparisons P value<0.0001, one-way repeated measure ANOVA, followed by Tukey’s multiple comparisons test).

pMKK4 was present in the cerebellum and both Tg and nTg hippocampus, and in both regions it is reduced by DLKi, demonstrating that pMKK4 is in part constitutively phosphorylated by DLK in that region. Interestingly, pMKK4 is also present and partially repressed by DLKi in the forebrain and hippocampus (Fig. 5E). Thus DLK has some baseline activity in the cerebellum, forebrain, and hippocampus, but only in the cerebellum does this activity culminate in extensive nuclear p-c-Jun under normal conditions.

### DLK scaffolding proteins are differentially expressed in cerebellum versus cortex and hippocampus

We reasoned that there must be differences in DLK signaling cofactors that explain why DLK activity leads to abundant nuclear p-c-Jun in the cerebellum but not in the forebrain or hippocampus in the absence of neuronal stress. We then examined our RNAseq data (see below) for regional disparities in expression of known JNK scaffolding genes. Of the genes examined, only one, *Sh3rf1* (POSH), was highly regionally enriched, with >10-fold greater expression in the hippocampus than in the cerebellum. We then compared other known cofactors by western blot. DLK expression was the same across brain regions, but JIP-1 was predominant in the cerebellum as previously reported (48), whereas JIP3, POSH, pMKK4, JNK1, and JNK3 were enriched in the hippocampus and forebrain. Thus the differential availability of signaling partners could determine whether nuclear p-c-Jun results from DLK activity.

**Fig. 6.**
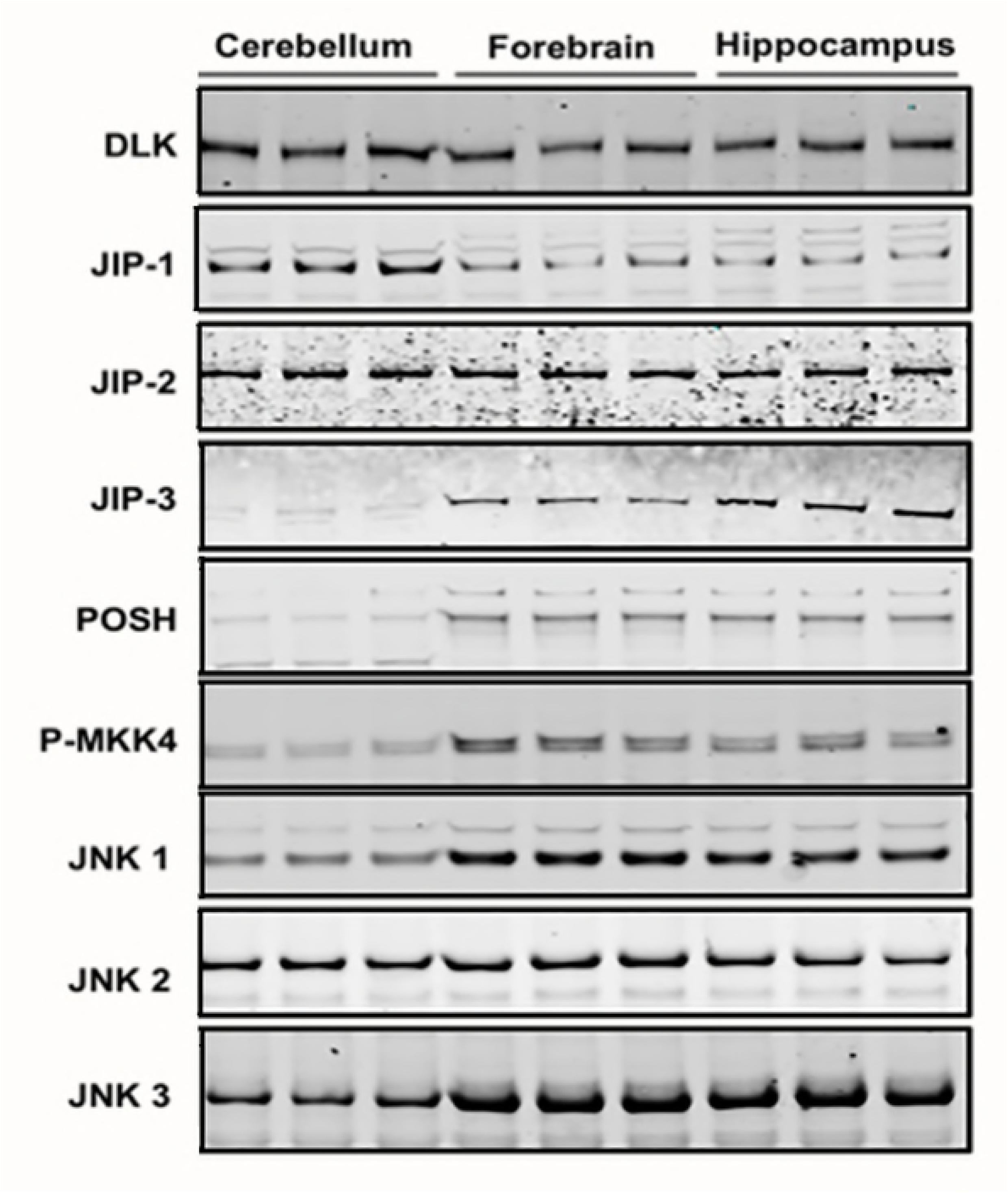
DLK has different scaffolding proteins that form the DLK/JNK signalsome in cerebellum versus hippocampus. Immunoblot analysis from lysates obtained from cerebellum, forebrain and hippocampus of 3-4 months old C57BL/6 mice.

### Downstream Consequences of DLK Activity are Regionally Distinct

Since DLK signals through activation of transcription factors such as p-c-Jun (49) changes in mRNA levels after DLK inhibition could identify differences in signaling output. We performed RNAseq of both hippocampus and cerebellum from nTg and Tg mice 50 hours after continuous inhibition by DLKi. ANOVA and post-hoc tests accounting for brain region, genotype, and treatment identified significant (p<0.05) differentially expressed genes in the DLKi treatment groups (Fig. 7A,B). In the hippocampus, roughly similar number of genes were upregulated as were downregulated by DLKi, with substantially more genes reaching significance in the Tg compared to nTg hippocampus, consistent with the elevated p-c-Jun in the disease model. In contrast, approximately 3000 genes were differentially expressed in the cerebellum and were 2-3-fold more likely to be downregulated, indicating that constitutive DLK activity in the cerebellum primarily induces gene expression (Fig. 7C). For reasons we do not yet understand, the Tg cerebellum was more sensitive to DLKi than the nTg cerebellum despite the absence of pathological tau in that brain region.

**Fig. 7.**
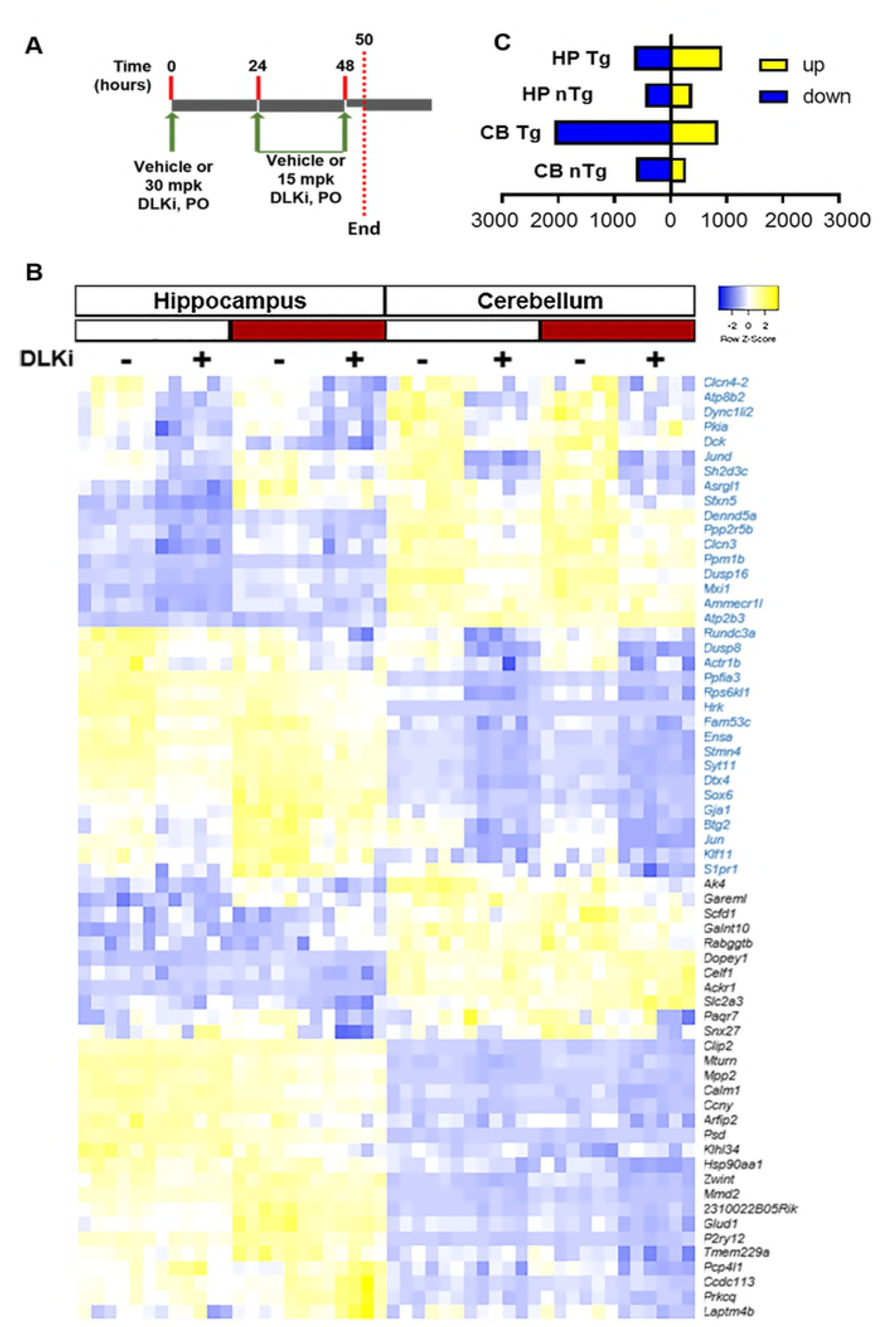
RNA seq analysis reveals a DLK-dependent gene signature in both cerebellum and hippocampus in nTg and Tg mice. **(A)** Schematic representation of study timeline. nTg (n=6) and Tg (n=6) mice were treated with vehicle or DLKi (30 mg/kg, qd, day 1 and 15 mg/kg, qd, days 2-3). The study was terminated approximately 2 hours after the last dose at maximal plasma concentrations. **(B)** Heatmap depicting the differentially expressed genes (p<0.05) due to DLK inhibition in which relative gene expression changes (blue = down; yellow = up) across genotype, treatment and brain region for each sample (n=6 samples/group). Upper genes listed (in blue text) depict genes significantly altered by the DLKi irrespective of genotype or brain region. Lower genes listed (in black text) depict genes altered by DLK inhibition in a genotype or regionally selective manner. **(C)** Number of differentially expressed genes (yellow = up, blue – down; p<0.05) due to DLK inhibition for each brain region*genotype is plotted.

Heatmap visualization showed that some genes were affected by DLKi in both brain regions while others showed regional dependence. Among the transcripts differentially expressed following DLKi treatment regardless of region are known DLK pathway components such as *Jun*, *Jund*, and the JNK phosphatases *Dusp8* and *Dusp16* (Fig 8A). We examined other genes consistently regulated by DLKi to identify potential novel pathway components. Four genes showed similar regulation patterns to these above known pathway members such as *Jun*. These include the mRNA for pro-apoptotic Bcl-2 relative Harakiri (*Hrk*); *Sh2d3c*, also known as SHEP-1, a scaffolding protein implicated in JNK and Cas signaling (50); *Stmn4*, a stathmin family member that induces microtubule destabilization (51); and *Rps6kl1*, encoding a potential kinase that has not been studied (Fig. 8B). Finally, we submitted the combined differentially expressed genes (Fig. 7B) to regulatory element analysis to identify putative transcription factors mediating the response to DLKi. Predicted Jun binding sites were the most significantly enriched (Fig. 8D), suggesting that p-c-Jun is a primary means by which DLK regulates transcription in these regions.

**Fig. 8.**
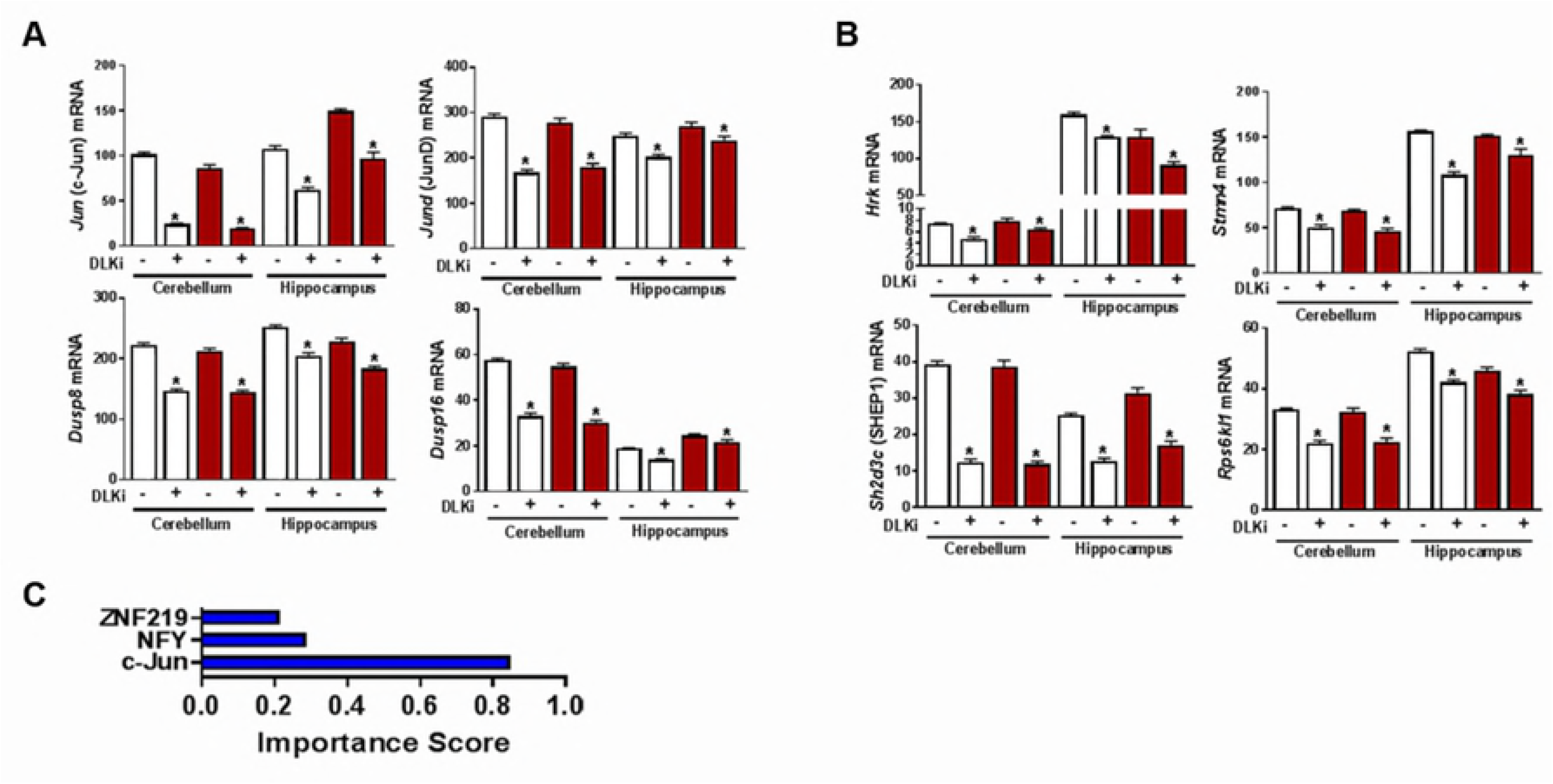
Pharmacological Inhibition of DLK decreases the expression of several genes within the. **(A)** MAPK pathway and **(B)** PERK pathway and **(C)** genes co-regulated with known DLK/JNK signaling factors. Data are plotted as mean mRNA expression levels (raw counts) + SEM (n=6) * indicates a significant change (p<0.05) as assessed via ANOVA of RNAseq data. **(D)** Promoter analysis using DIRE of the most significantly altered DLK-dependent genes (listed in (Fig. 7B)) indicates c-Jun is the most significantly represented (Importance score) transcriptional element.

### Pathway Analysis of *Jun* Co-regulated Genes

Next we mined the gene expression data to understand the differential function of DLK in the nTg cerebellum versus Tg hippocampus. Given that *Jun* mRNA induction is a surrogate for DLK-dependent nuclear p-c-Jun activity, a Pearson correlation analysis was performed to identify the genes that are co-regulated with *Jun* mRNA in both contexts. 688 and 1104 genes were significantly correlated with *Jun* mRNA (p<0.05) in nTg DLKi cerebellum and Tg DLKi hippocampal samples, respectively. KEGG pathway analysis found 7 pathways to be significantly altered in nTg DLKi cerebellum, mostly involving basic metabolism pathways (Fig. 9A), whereas 10 pathways showed significant enrichment in Tg DLKi hippocampus samples, most pathways indicating involvement in immune cell function (Fig. 9B). Thus constitutive DLK signaling in the cerebellum differs in functional output from the disease-induced DLK signaling in the Tg hippocampus.

**Fig. 9.**
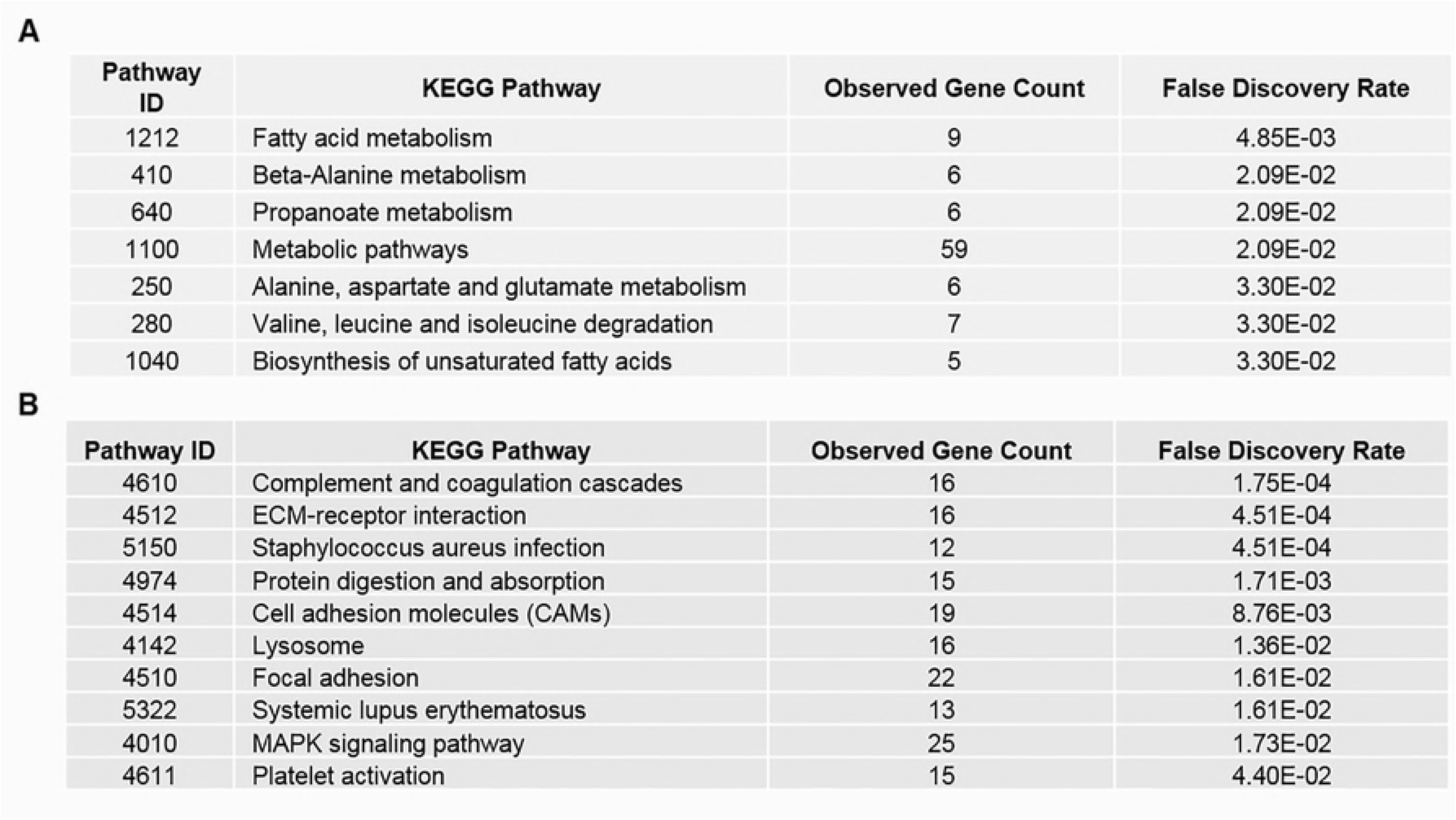
KEGG Pathway Analysis of genes most highly correlated with *Jun*. A Pearson correlation analysis of gene expression levels of all genes to *Jun* mRNA levels across either all nTg DLKi cerebellum samples or all Tg DLKi hippocampal samples was performed. KEGG pathways showing significant enrichment for significantly correlated (p<0.05) indicated significant enrichment (false discovery rate p<0.05) of **(A)** 7 pathways for nTg DLKi cerebellum samples, and **(B)** 10 pathways for Tg DLKi hippocampal samples.

## Discussion

The MAPK/JNK signaling pathway performs diverse, sometimes opposing functions, ranging from apoptosis to proliferation, depending on context. Here we illustrate that this diversity of function extends to DLK, which is not only a regulator of developmental processes and injury response, but also performs homeostatic functions likely related to synaptic function in the mature cerebellum. We observed robust nuclear p-c-Jun in cerebellar granule cells in adult mice, and by acutely blocking DLK with a highly selective inhibitor, p-c-Jun levels decreased rapidly. Given the highly selective nature of this DLKi, this decrease in p-c-Jun is likely due predominantly to DLK inhibition, although a minor contribution by LZK, which can also activate the JNK pathway, cannot be excluded.

This observation means that, in addition to mechanisms that activate DLK in injured and developing neurons, there must also be as-yet undefined processes that allow DLK to trigger p-c-Jun nuclear accumulation in cerebellar granule cells and/or that restrict c-Jun phosphorylation in other types of neurons. This mechanism does not appear to be driven by DLK stabilization, as has been observed in various peripheral nerve injury models (12, 18, 23-25, 51), because DLK levels were constant in all regions and conditions tested. DLK’s direct substrate pMKK4 is ~50% reduced by DLKi in both the hippocampus and cerebellum regardless of disease status, implying that DLK is constitutively active in both brain regions, and that regulation occurs further downstream. We suspect this regulation occurs at the assembly of the signaling complex, because DLK/JNK scaffolding proteins 1) interact with motor proteins to allow transport of the signaling complex to the perinuclear area and 2) can recruit negative regulatory factors such as phosphatases that limit signaling (32). Thus scaffolding of the DLK signalsome could be a key regulatory step. Support for this idea comes from the regional distribution of JIP-1, JIP-3, and POSH. JIP-1 is more abundant in the cerebellum, whereas JIP-3 and POSH are more abundant in the hippocampus, presumably leading to differences in DLK/JNK scaffolding and thus the outcome of DLK signaling. In dorsal root ganglion neurons, JIP-3, but not JIP-1, is essential for DLK-mediated apoptosis (52), highlighting the functional differences associated with the availability of specific scaffolding protein, and raising the possibility that the lower JIP-3 in the cerebellum promotes physiological DLK signaling. The strikingly higher levels of POSH in the hippocampus could also be functionally important. POSH forms a complex with JIP-1 (53) and like JIP-1, is required for injury-induced hippocampal apoptosis (54, 55). Perhaps JIP-3 and putative JIP-1/POSH heterodimers limit DLK signaling in healthy hippocampal neurons but permit it following injury. Future studies will be required to test this hypothesis.

Several genes were identified as commonly repressed in all conditions by DLKi; these genes include *Jun*, *Jund*, the JNK phosphatases *Dusp8* and *Dusp16*, and other components of the DLK/JNK signaling pathway. Reasoning that genes co-regulated with these known factors might be novel components of DLK signaling, we searched for transcripts that were also consistently down-regulated by DLKi and identified four genes of interest. Of note is Harakiri (*Hrk*), a c-Jun direct target gene that was previously shown to be DLK-induced (5, 56) and is required for neuronal death after trophic factor withdrawal or axotomy in vitro (57). *Hrk* mRNA is >20-fold higher in the hippocampus than the cerebellum (Fig. 6e), suggesting that it could contribute to regional differences in DLK signaling relating to injury responses.

The comparative transcriptomics data also show that DLK signaling can have significantly different outcomes depending on context, not only regionally, but under normal physiological conditions or disease. Pathway analyses of the cerebellum data showed that the genes co-regulated with Jun mRNA were enriched in basic metabolic pathways, suggesting a DLK-mediated link between synaptic function and metabolism. These data are consistent with the strong correlation we observed between p-c-jun and synaptophysin, and work in mice and invertebrates demonstrating a role for DLK in regulating synaptic strength (16). No gross changes in motor function or ambulatory behavior after long-term conditional DLK knockdown has been reported, thus the impact on DLK signaling on cerebellar physiology remains unknown. Detailed electrophysiological experiments in the DLK adult somatic knockout mouse will be important to reveal this function.

Compared to the cerebellum, the role of DLK under normal physiological conditions in the hippocampus appears minimal, as DLK inhibition led to very few gene expression changes. When high levels of pathological tau in the hippocampus cause DLK-dependent nuclear p-c-Jun, far more genes were differentially expressed. Here the *Jun* co-regulated gene set maps functionally to pathways involved in immune cell function such as complement pathway, which is involved in developmental and pathological synapse elimination (58). Future studies will determine if DLK-dependent crosstalk between injured neurons and microglia, which produce complement, contribute to synaptic loss in injury and neurodegeneration.

## Acknowledgments

We thank the Robert A and Renee E. Belfer Family Foundation and the many donors to the University of Texas MD Anderson Cancer Center who supported this work, as well as the Alzheimer’s Disease Drug Discovery Foundation (Grant 20160903, WJR). We also thank Quanyun Xu for the bioanalytical analyses and Dr. Suparna Sanyal for her contributions to the manuscript.

---

**Fig. S1 Validation of DLK antibodies. (A)** Immunoblot analysis of DLK using DLK antibodies from two vendors in whole brain lysates from DLK WT, DLK (+/-) and DLK (-/-) mice **(B)** Immunoblot analysis of immunoprecipitated DLK from adult wildtype C57Bl6 mice brain (IP with DLK antibody, Thermo Scientific; immunoblot with DLK antibody, NeuroMab).

## References

1. Conforti L, Gilley J, Coleman MP. Wallerian degeneration: an emerging axon death pathway linking injury and disease. Nat Rev Neurosci. 2014;15(6):394–409.

2. Miller BR, Press C, Daniels RW, Sasaki Y, Milbrandt J, DiAntonio A. A dual leucine kinase-dependent axon self-destruction program promotes Wallerian degeneration. Nat Neurosci. 2009;12(4):387–9.

3. Tedeschi A, Bradke F. The DLK signalling pathway--a double-edged sword in neural development and regeneration. EMBO Rep. 2013;14(7):605–14.

4. Hammarlund M, Nix P, Hauth L, Jorgensen EM, Bastiani M. Axon regeneration requires a conserved MAP kinase pathway. Science. 2009;323(5915):802–6.

5. Watkins TA, Wang B, Huntwork-Rodriguez S, Yang J, Jiang Z, Eastham-Anderson J, et al. DLK initiates a transcriptional program that couples apoptotic and regenerative responses to axonal injury. Proc Natl Acad Sci U S A. 2013;110(10):4039–44.

6. Welsbie DS, Yang Z, Ge Y, Mitchell KL, Zhou X, Martin SE, et al. Functional genomic screening identifies dual leucine zipper kinase as a key mediator of retinal ganglion cell death. Proc Natl Acad Sci U S A. 2013;110(10):4045–50.

7. Xiong X, Collins CA. A conditioning lesion protects axons from degeneration via the Wallenda/DLK MAP kinase signaling cascade. J Neurosci. 2012;32(2):610–5.

8. Xiong X, Wang X, Ewanek R, Bhat P, Diantonio A, Collins CA. Protein turnover of the Wallenda/DLK kinase regulates a retrograde response to axonal injury. J Cell Biol. 2010;191(1):211–23.

9. Yan D, Wu Z, Chisholm AD, Jin Y. The DLK-1 kinase promotes mRNA stability and local translation in C. elegans synapses and axon regeneration. Cell. 2009;138(5):1005–18.

10. Hirai S, Cui DF, Miyata T, Ogawa M, Kiyonari H, Suda Y, et al. The c-Jun N-terminal kinase activator dual leucine zipper kinase regulates axon growth and neuronal migration in the developing cerebral cortex. J Neurosci. 2006;26(46):11992–2002.

11. Hirai S, Banba Y, Satake T, Ohno S. Axon formation in neocortical neurons depends on stage-specific regulation of microtubule stability by the dual leucine zipper kinase-c-Jun N-terminal kinase pathway. J Neurosci. 2011;31(17):6468–80.

12. Nakata K, Abrams B, Grill B, Goncharov A, Huang X, Chisholm AD, et al. Regulation of a DLK-1 and p38 MAP kinase pathway by the ubiquitin ligase RPM-1 is required for presynaptic development. Cell. 2005;120(3):407–20.

13. Itoh A, Horiuchi M, Wakayama K, Xu J, Bannerman P, Pleasure D, et al. ZPK/DLK, a mitogen-activated protein kinase kinase kinase, is a critical mediator of programmed cell death of motoneurons. J Neurosci. 2011;31(20):7223–8.

14. Klinedinst S, Wang X, Xiong X, Haenfler JM, Collins CA. Independent pathways downstream of the Wnd/DLK MAPKKK regulate synaptic structure, axonal transport, and injury signaling. J Neurosci. 2013;33(31):12764–78.

15. Crawley O, Giles AC, Desbois M, Kashyap S, Birnbaum R, Grill B. A MIG-15/JNK-1 MAP kinase cascade opposes RPM-1 signaling in synapse formation and learning. PLoS Genet. 2017;13(12):e1007095.

16. Pozniak CD, Sengupta Ghosh A, Gogineni A, Hanson JE, Lee SH, Larson JL, et al. Dual leucine zipper kinase is required for excitotoxicity-induced neuronal degeneration. J Exp Med. 2013;210(12):2553–67.

17. Le Pichon CE, Meilandt WJ, Dominguez S, Solanoy H, Lin H, Ngu H, et al. Loss of dual leucine zipper kinase signaling is protective in animal models of neurodegenerative disease. Sci Transl Med. 2017;9(403).

18. Lewcock JW, Genoud N, Lettieri K, Pfaff SL. The ubiquitin ligase Phr1 regulates axon outgrowth through modulation of microtubule dynamics. Neuron. 2007;56(4):604–20.

19. Yan D, Jin Y. Regulation of DLK-1 kinase activity by calcium-mediated dissociation from an inhibitory isoform. Neuron. 2012;76(3):534–48.

20. Holland SM, Collura KM, Ketschek A, Noma K, Ferguson TA, Jin Y, et al. Palmitoylation controls DLK localization, interactions and activity to ensure effective axonal injury signaling. Proc Natl Acad Sci U S A. 2016;113(3):763–8.

21. Li J, Zhang YV, Asghari Adib E, Stanchev DT, Xiong X, Klinedinst S, et al. Restraint of presynaptic protein levels by Wnd/DLK signaling mediates synaptic defects associated with the kinesin-3 motor Unc-104. Elife. 2017;6.

22. Valakh V, Frey E, Babetto E, Walker LJ, DiAntonio A. Cytoskeletal disruption activates the DLK/JNK pathway, which promotes axonal regeneration and mimics a preconditioning injury. Neurobiol Dis. 2015;77:13–25.

23. Collins CA, Wairkar YP, Johnson SL, DiAntonio A. Highwire restrains synaptic growth by attenuating a MAP kinase signal. Neuron. 2006;51(1):57–69.

24. Huntwork-Rodriguez S, Wang B, Watkins T, Ghosh AS, Pozniak CD, Bustos D, et al. JNK-mediated phosphorylation of DLK suppresses its ubiquitination to promote neuronal apoptosis. J Cell Biol. 2013;202(5):747–63.

25. Nix P, Hisamoto N, Matsumoto K, Bastiani M. Axon regeneration requires coordinate activation of p38 and JNK MAPK pathways. Proc Natl Acad Sci U S A. 2011;108(26):10738–43.

26. Larhammar M, Huntwork-Rodriguez S, Rudhard Y, Sengupta-Ghosh A, Lewcock JW. The Ste20 Family Kinases MAP4K4, MINK1, and TNIK Converge to Regulate Stress-Induced JNK Signaling in Neurons. J Neurosci. 2017;37(46):11074–84.

27. Xu Z, Maroney AC, Dobrzanski P, Kukekov NV, Greene LA. The MLK family mediates c-Jun N-terminal kinase activation in neuronal apoptosis. Mol Cell Biol. 2001;21(14):4713–24.

28. Mata M, Merritt SE, Fan G, Yu GG, Holzman LB. Characterization of dual leucine zipper-bearing kinase, a mixed lineage kinase present in synaptic terminals whose phosphorylation state is regulated by membrane depolarization via calcineurin. J Biol Chem. 1996;271(28):16888–96.

29. Asaoka Y, Nishina H. Diverse physiological functions of MKK4 and MKK7 during early embryogenesis. J Biochem. 2010;148(4):393–401.

30. Bode AM, Dong Z. The functional contrariety of JNK. Mol Carcinog. 2007;46(8):591–8.

31. Lee CM, Onesime D, Reddy CD, Dhanasekaran N, Reddy EP. JLP: A scaffolding protein that tethers JNK/p38MAPK signaling modules and transcription factors. Proc Natl Acad Sci U S A. 2002;99(22):14189–94.

32. Mooney LM, Whitmarsh AJ. Docking interactions in the c-Jun N-terminal kinase pathway. J Biol Chem. 2004;279(12):11843–52.

33. Whitmarsh AJ. The JIP family of MAPK scaffold proteins. Biochem Soc Trans. 2006;34(Pt 5):828–32.

34. Xu Z, Kukekov NV, Greene LA. POSH acts as a scaffold for a multiprotein complex that mediates JNK activation in apoptosis. EMBO J. 2003;22(2):252–61.

35. Coffey ET. Nuclear and cytosolic JNK signalling in neurons. Nat Rev Neurosci. 2014;15(5):285–99.

36. Herdegen T, Skene P, Bahr M. The c-Jun transcription factor--bipotential mediator of neuronal death, survival and regeneration. Trends Neurosci. 1997;20(5):227–31.

37. Dobin A, Davis CA, Schlesinger F, Drenkow J, Zaleski C, Jha S, et al. STAR: ultrafast universal RNA-seq aligner. Bioinformatics. 2013;29(1):15–21.

38. Li H, Handsaker B, Wysoker A, Fennell T, Ruan J, Homer N, et al. The Sequence Alignment/Map format and SAMtools. Bioinformatics. 2009;25(16):2078–9.

39. Anders S, Pyl PT, Huber W. HTSeq--a Python framework to work with high-throughput sequencing data. Bioinformatics. 2015;31(2):166–9.

40. Babicki S, Arndt D, Marcu A, Liang Y, Grant JR, Maciejewski A, et al. Heatmapper: web-enabled heat mapping for all. Nucleic Acids Res. 2016;44(W1):W147–53.

41. Hamby ME, Coppola G, Ao Y, Geschwind DH, Khakh BS, Sofroniew MV. Inflammatory mediators alter the astrocyte transcriptome and calcium signaling elicited by multiple G-protein-coupled receptors. J Neurosci. 2012;32(42):14489–510.

42. Gotea V, Ovcharenko I. DiRE: identifying distant regulatory elements of co-expressed genes. Nucleic Acids Res. 2008;36(Web Server issue):W133–9.

43. Szklarczyk D, Franceschini A, Wyder S, Forslund K, Heller D, Huerta-Cepas J, et al. STRING v10: protein-protein interaction networks, integrated over the tree of life. Nucleic Acids Res. 2015;43(Database issue):D447–52.

44. Suenaga J, Cui DF, Yamamoto I, Ohno S, Hirai S. Developmental changes in the expression pattern of the JNK activator kinase MUK/DLK/ZPK and active JNK in the mouse cerebellum. Cell Tissue Res. 2006;325(1):189–95.

45. Yuan Z, Gong S, Luo J, Zheng Z, Song B, Ma S, et al. Opposing roles for ATF2 and c-Fos in c-Jun-mediated neuronal apoptosis. Mol Cell Biol. 2009;29(9):2431–42.

46. Patricelli MP, Nomanbhoy TK, Wu J, Brown H, Zhou D, Zhang J, et al. In situ kinase profiling reveals functionally relevant properties of native kinases. Chem Biol. 2011;18(6):699–710.

47. Santacruz K, Lewis J, Spires T, Paulson J, Kotilinek L, Ingelsson M, et al. Tau suppression in a neurodegenerative mouse model improves memory function. Science. 2005;309(5733):476–81.

48. Pellet JB, Haefliger JA, Staple JK, Widmann C, Welker E, Hirling H, et al. Spatial, temporal and subcellular localization of islet-brain 1 (IB1), a homologue of JIP-1, in mouse brain. Eur J Neurosci. 2000;12(2):621–32.

49. Larhammar M, Huntwork-Rodriguez S, Jiang Z, Solanoy H, Sengupta Ghosh A, Wang B, et al. Dual leucine zipper kinase-dependent PERK activation contributes to neuronal degeneration following insult. Elife. 2017;6.

50. Sakakibara A, Hattori S. Chat, a Cas/HEF1-associated adaptor protein that integrates multiple signaling pathways. J Biol Chem. 2000;275(9):6404–10.

51. Chauvin S, Sobel A. Neuronal stathmins: a family of phosphoproteins cooperating for neuronal development, plasticity and regeneration. Prog Neurobiol. 2015;126:1–18.

52. Ghosh AS, Wang B, Pozniak CD, Chen M, Watts RJ, Lewcock JW. DLK induces developmental neuronal degeneration via selective regulation of proapoptotic JNK activity. J Cell Biol. 2011;194(5):751–64.

53. Cunningham CA, Knudson KM, Peng BJ, Teixeiro E, Daniels MA. The POSH/JIP-1 scaffold network regulates TCR-mediated JNK1 signals and effector function in CD8(+) T cells. Eur J Immunol. 2013;43(12):3361–71.

54. Whitmarsh AJ, Kuan CY, Kennedy NJ, Kelkar N, Haydar TF, Mordes JP, et al. Requirement of the JIP1 scaffold protein for stress-induced JNK activation. Genes Dev. 2001;15(18):2421–32.

55. Zhang QG, Wang RM, Yin XH, Pan J, Xu TL, Zhang GY. Knock-down of POSH expression is neuroprotective through down-regulating activation of the MLK3-MKK4-JNK pathway following cerebral ischaemia in the rat hippocampal CA1 subfield. J Neurochem. 2005;95(3):784–95.

56. Towers E, Gilley J, Randall R, Hughes R, Kristiansen M, Ham J. The proapoptotic dp5 gene is a direct target of the MLK-JNK-c-Jun pathway in sympathetic neurons. Nucleic Acids Res. 2009;37(9):3044–60.

57. Imaizumi K, Benito A, Kiryu-Seo S, Gonzalez V, Inohara N, Lieberman AP, et al. Critical role for DP5/Harakiri, a Bcl-2 homology domain 3-only Bcl-2 family member, in axotomy-induced neuronal cell death. J Neurosci. 2004;24(15):3721–5.

58. Hong S, Beja-Glasser VF, Nfonoyim BM, Frouin A, Li S, Ramakrishnan S, et al. Complement and microglia mediate early synapse loss in Alzheimer mouse models. Science. 2016;352(6286):712–6.

59. Andrews, S. (2010). FastQC: A quality control tool for High Throughput Sequence Data. Retrieved from http://www.bioinformatics.babraham.ac.uk/projects/fastqc

60. Martin, Marcel. Cutadapt removes adapter sequences from high-throughput sequencing reads. EMBnet.journal, [S.l.], v. 17, n. 1, p. pp. 10–12, may 2011. ISSN 2226-6089.

